# Coordinated action of RTBV and RTSV proteins suppress host RNA silencing machinery

**DOI:** 10.1101/2021.01.19.427099

**Authors:** Abhishek Anand, Malathi Pinninti, Anita Tripathi, Satendra Kumar Mangrauthia, Neeti Sanan-Mishra

## Abstract

RNA silencing is as an adaptive immune response in plants that limits accumulation or spread of invading viruses. Successful virus infection entails countering the RNA silencing for efficient replication and systemic spread in the host. The viruses encode proteins having the ability to suppress or block the host silencing mechanism, resulting in severe pathogenic symptoms and diseases. Tungro virus disease caused by a complex of two viruses provides an excellent system to understand these host and virus interactions during infection. It is known that *Rice tungro bacilliform virus* (RTBV) is the major determinant of the disease while *Rice tungro spherical virus* (RTSV) accentuates the symptoms. This study brings to focus the important role of RTBV ORF-IV in Tungro disease manifestation, by acting as both the victim and silencer of the RNA silencing pathway. The ORF-IV is a weak suppressor of the S-PTGS or pre-established stable silencing but its suppression activity is augmented in the presence of RTSV proteins. The RTBV and RTSV proteins interact to suppress localized silencing as well as spread of silencing, in the host plants.

## Introduction

Viruses represent the most invasive group of pathogens that are harmful for any host system they infect. Viral diseases in cultivated crops negatively affect plant morphology and physiology, thereby resulting in huge losses. However, resistance against viruses was observed as early as 1929 followed by observations of the phenomenon popularly known as “cross protection”. In this a plant infected with one virus showed resistance to the same or closely related virus on successive infection (McKinney, 1929). Subsequently it was shown that the plants having disease symptoms on virus infection quickly recovered and the new tissues emerged without any symptoms (Lindbo et al., 1993). This served as the early basis for engineering plants for viral resistance by integrating the viral genetic material into the plant genome and was termed as “pathogen derived resistance” (PDR) (Lindbo, 1992; Van der Vlugt and Goldbach 1992; Wilson, 1993; Beachy, 1997; Ratcliff et al., 1997). It was later explained that the cross protection and the recovery by PDR were associated with small RNA mediated gene silencing at molecular level. This phenomenon emerged as one of the most well-developed, robust and potent defense strategies employed by plants and other eukaryotes (Ruiz et al., 1998; Baulcombe, 1999; Mansoor et al., 2006; Ding, 2010; Vaucheret et al., 2001; Baulcombe, 2004).

The small RNA mediated gene silencing is a sequence specific regulatory phenomenon resulting in mRNA degradation or translation inhibition at the post-transcriptional level and DNA methylation at transcriptional level (Agrawal et al., 2003; Sanan-Mishra and Mukherjee 2007; Pelaez and Sanchez, 2013; Sanan-Mishra et al., 2013). The process involves generation of 21 to 24 nt long double stranded RNA intermediates by action of specific Dicer like (DCL) molecules that guide the silencing through Argonaute (AGO) containing protein complexes (Peragine et al., 2004; Sanan-Mishra et al., 2013). Depending on the mechanisms of biogenesis and function, the small RNAs are classified into microRNAs (miRNAs) and short interfering RNAs (siRNAs).

For successfully invading the hosts, viruses have co-evolved tools to effectively suppress the components of host silencing (Anandalakshmi et al., 1998; Anand et al., 2013). These viral encoded suppressor molecules have been recognized as pathogenicity determinants as they have the ability to counteract antiviral silencing and play an important role in virulence (Qu and Morris 2005; Omarov and Scholthof 2012; Kumar et al. 2015). The suppression activity has evolved independently such that existing viral proteins performing certain vital functions have gained an additional role of suppressing RNA silencing (Carrington et al., 2001; Li and Ding, 2001; Silhavy et al., 2002). Potyviral HC-Pro protein is the first and best described suppressor of host RNA silencing (Anandalakshmi et al., 1998). It is a multifunctional protein responsible for viral genome amplification, polyprotein processing, insect transmission and long distance movement (Kasschau et al., 1997). It also acts as a broad range pathogenicity enhancer by suppressing host silencing machinery (Pruss et al., 1997). The suppressor proteins do not share any co-evolutionary patterns among different viruses and there is very little chance of any significant homology or sequence similarity between them. Moreover, they have unique functional domains and properties, resulting in distinctive mechanism of interacting with the components of host RNA silencing machinery (Carrington et al., 2001; Li and Ding, 2001; Silhavy et al., 2002). All these facts add up to complicate their identification and classification. However their ability to suppress RNA silencing, regardless of system or sequence, has been exploited in excellent assays to characterize the suppressors. Number of *in planta* assays are available based on the suppression of RNA silencing of a reporter transgene (Palauqui et al., 1997; Voinnet and Baulcombe, 1997; Voinnet, 2005; Karjee et al., 2008). The principle behind these assays is that the silencing of a reporter transgene would be reversed by the effect of the suppressor when it is introduced and expressed in the system.

Tungro disease of rice is considered as one of the most damaging viral diseases of rice prevalent in South and Southeast Asia and accounts for huge economic loss (Azzam and Chancellor, 2002). It is caused by a complex of two viruses, transmitted together by the green leafhoppers (GLH) - *Nephotettix virescens* and *N. nigropictus* (Cabauatan and Hibino, 1985). The *Rice tungro spherical virus* (RTSV), a positive single strand RNA virus in the Secoviridae family (Sanfaçon et al., 2009), is required for transmission of disease (**Fig S1**). Single infection by RTSV does not produce any detectable symptoms of the disease. Its genome consists of 12433-nt RNA with a poly-A tail at the 3’ end that codes for a large polyprotein, which is cleaved to form three mature capsid proteins and other viral proteins (Shen et al., 1993; Kunii et al., 2004). *Rice tungro bacilliform virus* (RTBV), a double strand DNA pararetrovirus virus in the Caulimoviridae family (**Fig S1**) is considered as the major cause for the manifestation of the disease (Hibino et al., 1978; Jones et al., 1991; Hull, 1996; Mangrauthia et al, 2012). RTBV genome contains four open reading frames (ORFs). Functions of ORF-I and ORF-II are yet not defined. The ORF-III codes for a large polyprotein, which is processed in to a movement protein (MP), a coat protein (CP), an aspartate protease (AP) and a replicase proteins (Hay et al., 1991). The ORF-IV transcript undergoes splicing to generate a protein that has a sequence motif similar to leucine zipper although it lacks the DNA binding region. Though RTBV is the main determinant of disease symptoms, the presence of RTSV accentuates the symptoms (Hull, 1996; Borah et al., 2013). Stunting, yellow orange discoloration of leaves and twisting of leaf tips are the major symptoms in plants infected with Tungro virus (Hibino et al., 1978; Dasgupta et al., 1991; Srilatha et al, 2019).

It was demonstrated that transgenic rice plants expressing DNA encoding ORF-IV of RTBV, both in sense as well as in anti-sense orientation were effective for controlling RTBV infection (Tyagi et al., 2008). This study suggested that there is an involvement of small RNA molecules in providing resistance towards the disease. It was also reported that the rice silencing machinery responds to RTBV infection by generating typical viral siRNAs that are potentially associated with multiple AGOs in active RISC to direct silencing of the viral genes. However, RTBV appears to evade the repressive action of viral siRNAs by restricting their production to the non-coding region. It was also demonstrated that the protein encoded by ORF-IV of RTBV suppressed cell autonomous silencing (Rajeswaran et al., 2014).

In this work we show that complexity of disease is driven by combined suppressor action of two viral components of the tungro system. The ORF-IV protein of RTBV acts as a weak suppressor of RNA silencing but its suppression activity is augmented in the presence of the RTSV proteins. RTSV components might also have a possible role in suppressing the cell to cell spread of silencing. The suppressor activities were assayed by utilizing an *in-planta* assay based on the reversal of S-PTGS or pre-established stable GFP silencing (Karjee et al., 2008). The recognition of RTBV ORF-IV as a target and suppressor of the RNA silencing pathway has identified a key target that can be exploited for boosting host plant resistance against tungro virus infection.

## Materials and methods

### Plant material

Rice leaf tissues were collected from healthy and tungro infected plants grown in controlled conditions at ICAR-IIRR, Hyderabad. To obtain tungro infected rice tissue, 15 days old plants of susceptible cultivar, Taichung Native 1 (TN1) were inoculated with Hyderabad isolate of tungro virus using insect mediated virus transmission (Srilatha et al., 2019). The insect vectors, green leafhoppers (GLH), were fed on tungro disease infected plants for 24h (to acquire the virus particles) before releasing them onto healthy plants for 6h. The disease symptoms were clearly observed after 15 day of virus inoculation. The presence of virus was also confirmed by PCR using the methods described previously (Mangrauthia et al., 2010; Malathi and Mangrauthia, 2013). Thirty days old plants showing disease symptoms and virus presence were used for cDNA synthesis and small RNA library preparation.

### Small RNA library sequencing and computational analysis

Samples from three different biological replicates were pooled and used for library preparation as per manufacture’s (Illumina) protocol. The libraries thus made were used for deep sequencing on GAII sequencer (Illumina). This generated approx. 11M tags per library and data was delivered as sequences of 33 or 35 bases in length along with the base quality scores and read counts. The data has been deposited with NCBI GEO database (SAMN17245506). The obtained raw sequence reads were processed computationally using in house developed pipelines.

### Plasmid constructs

The individual ORFs of RTBV and RTSV were amplified from tungro infected plants and cloned into pGEMTeasy vectors. The inserts were moved into binary vector pBI121 under the CaMV 35S promoter and NOS terminator. The transformants were analyzed by restriction analysis, PCR and plasmid DNA sequencing. The recombinant plasmids (pBI121-ORF-IV, pBI121-ORF-I, pBI121-ORF-II, pBI121-ORF-Coat protein 3, pBI121-ORF-Polymerase and pBI121-ORF-protease) were mobilized into *Agrobacterium tumefaciens* strains LBA4404 cells.

### Agro-infiltration

Agrobacterium-mediated reversal of GFP expression in GFP silenced *Nicotiana tabacum* L. cv. Xanthi leaves (Karjee et al., 2008) was achieved through pressure infiltration, as described previously (Sparkes et al., 2006; Karjee et al., 2008; Das and Sana-Mishra, 2014). Briefly, Agrobacterium culture was grown until reaching an optical density of 1.0 at 600 nm (OD_600_). The culture was treated with 200 μM acetosyringone for 1h prior to infiltration. The homogenous culture mixture was infiltrated in the young leaves with the help of needleless syringe by generating a vacuum with the help of finger on the dorsal side of the leaf and mouth of the syringe on the ventral side.

### cDNA synthesis

First strand cDNA were prepared in 20 μl reactions from total RNA isolated from the infiltrated leaf tissues, using 50 U of Super Script TM II reverse transcriptase (Invitrogen) and random hexamers according to the manufacturers’ protocol. The first strand cDNA of total RNA were subjected to DNaseI treatment for 30 min.

### Reverse Transcriptase Polymerase Chain Reaction (RT-PCR)

The semi-quantitative RT-PCR, for the amplification of GFP and suppressor genes (RTBV ORF-IV, RTBV ORF I and RTSV ORF-protease) were carried out using gene specific primers at initial sample denaturation at 95°C for 5 min followed by 25 cycles of strand separation at 94°C for 1 min, annealing at 56°C for 30 s and extension at 72°C for 30 s. The program was extended for 7 min at 72°C. The tobacco 18S gene was used as a constitutive internal standard to evaluate cDNA content. The amplification products were analyzed on 0.8% agarose gel. The band intensities were quantified using Alpha Imager Imaging System.

### Northern blot analysis

Total RNA was extracted from infiltrated regions of the leaves using the guanidium thiocynate extraction method (Sambrook et al., 1989). 30 μg of total RNA from each plant sample was resolved on a formamide agarose gel, to check for GFP transcript and small RNA. The RNA was transferred on Hybond N^+^ membranes (Amersham Pharmacia). Digoxigenin-11-dUTP (DIG) labeled DNA probes were generated using via PCR labelling using PCR DIG Probe Synthesis kit (Roche, catalogue no. 11636090910). 10 ng of purified plasmid DNA containing full length GFP DNA was used as PCR template and amplified with GFP forward (TCAAGGACGACGGGAACTACAAG) and reverse (GTGGTGGTGGCTAGCTTTGTA) primers. Northern blot analysis was performed at 50°C using protocol provided in the DIG Application Manual for Filter Hybridization (Roche). Following the hybridization, the blot was washed at 55°C and probe detection was performed according to the manufacturers’ protocol using the DIG luminescent detection kit (Roche). The intensity of individual bands was measured with respect to the background and integrated density value (IDV) was calculated using Alphaimager. Each experiment was performed in triplicates and average values were used for plotting.

### Yeast two hybrid analysis

Coding sequences of ORF-IV of RTBV and CP1, CP2, CP3, P1, polymease and protease of RTSV were cloned into pGBKT7 and pGADT7 vectors, respectively. Yeast two hybrid assays were performed using the Matchmaker Gold Yeast Two-Hybrid System (Clontech). The standard protocol given in the kit was followed for Y2H assay. The yeast growth media (YPD medium, Clontech), SD growth medium w/o Ade^−^His^−^Leu^−^Trp^−^ (Himedia) and supplements media (Clontech) were used for the Y2H experiment.

## Results

### 1. ORF-IV of RTBV is a potential hotspot for siRNA generation

The small RNA sequencing data sets from tungro infected rice leaves (TL) were analyzed using in house developed pipelines to predict and map the siRNAs to RTBV genome. The small RNA reads were aligned to the RTBV genome sequence (Accession no. NC_001914.1) using Bowtie tool, allowing for only one mismatch at either first or last nucleotide. The TL small RNA libraries contained a total number of 2885141 reads, representing 554305 unique sequences. Read length distribution analysis showed that the 21-nt small RNAs, majorly representing the canonical miRNAs and siRNAs, are highest in number indicating a large diversity in their sequences (**Fig. 1A**). The first nucleotide of these molecules were either A or T in 68% of molecules (**Fig. 1A**), as shown in previous studies of rice siRNAs (Song et al., 2012; Wu et al., 2010; Wei et al., 2014; Rajeswaran et al., 2014).

**Figure 1.**
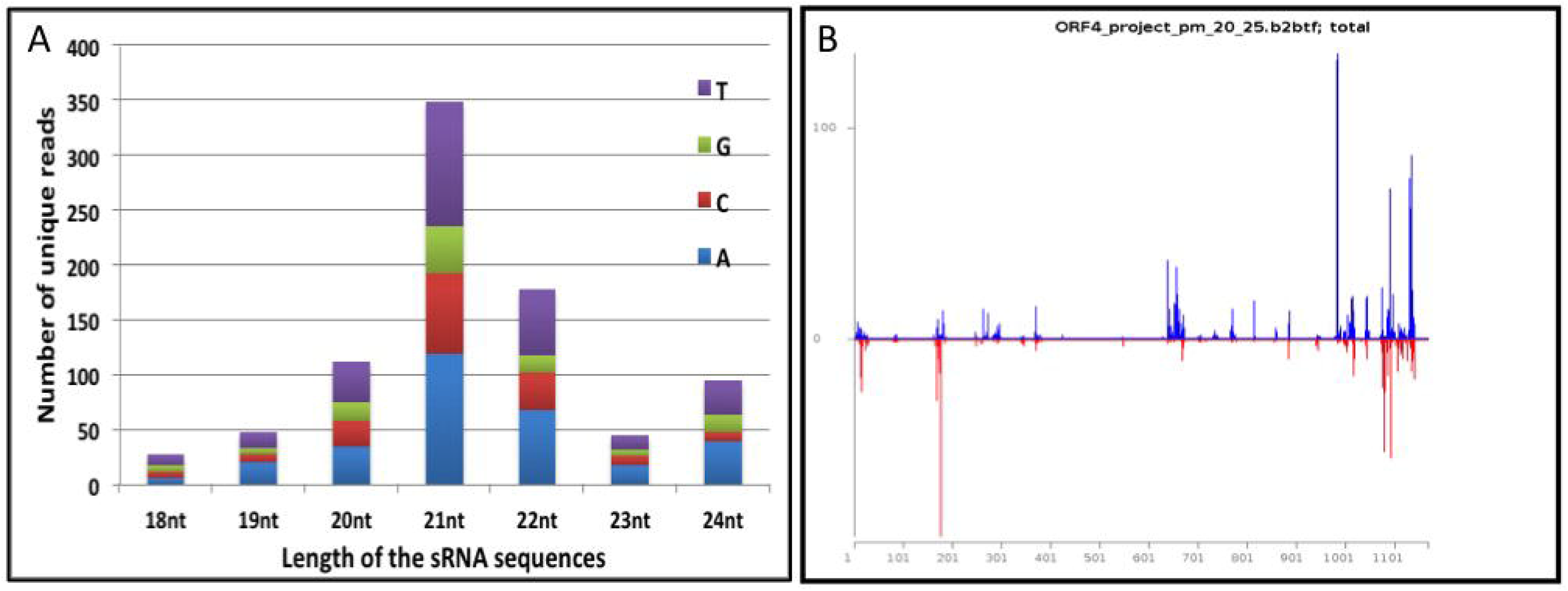
Size distribution and mapping of small RNA reads obtained from tungro infected data sets. **(A)** Length distribution of the small RNA sequences showing the distribution of first nucleotide in the molecules (**B**) Plot of sRNA molecules mapping to ORF-IV of RTBV. The siRNA mapping to positive strand of the ORF are represented by blue lines while those mapping to the than on negative strand are represented by red lines

The analysis identified 1690 unique reads aligned at different positions on both the strands of the RTBV genome. It was observed that ~50% (850) unique reads aligned at different positions distributed over most of the ORF-IV region although majority of siRNAs were clustered towards 3’ end. The abundance of siRNA biasing could also be observed on the positive strand of the ORF than on negative strand (**Fig. 1B**). Analysis of the mapping pattern and abundance of the putative siRNAs on the RTBV genome showed that ORF-IV acted as a probable hotspot for small RNA generation (**Fig. 1B**) in the infected leaf tissues.

### 2. ORF-IV can suppress pre-established RNA silencing

Leaves of stably silent GFP tobacco lines (Karjee et al., 2008) were infiltrated on one side of the midrib with Agrobacterium cultures containing the individual ORFs of RTBV (ORF-I, ORF-II and ORF-IV) and RTSV (protease, polymerase and coat protein 3). Leaves were collected at 3, 5 and 7 days post infiltration (dpi) and scanned under UV-light to observe for GFP fluorescence, as a proof for reversal of silencing by the infiltrated suppressor. The RTBV ORF-I, ORF-II and the RTSV ORFs did not show a clear suppressor activity in individual infiltrations (data not shown). In regions infiltrated with RTBV ORF-IV, low level of fluorescence was observed in leaves at 3dpi while significant fluorescence could be observed at 5dpi and this dipped again by 7dpi (**Fig. 2A**). No fluorescence was detected in the regions infiltrated with empty vector that served as control. This showed that ORF-IV acts as a weak suppressor of the S-PTGS or pre-established stable silencing.

**Figure 2.**
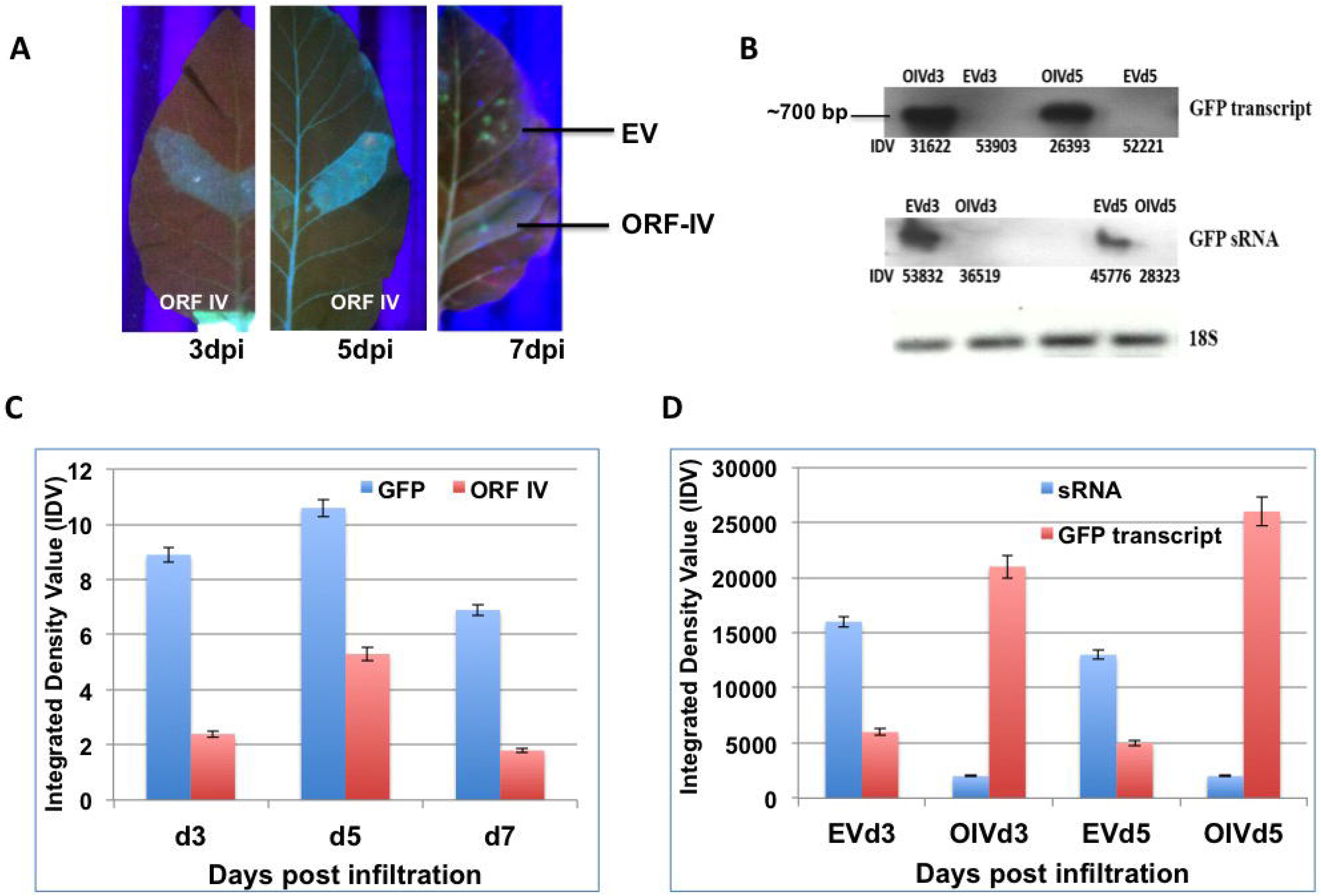
Reversal of GFP silencing by RTBV ORF-IV and its molecular analysis. (**A**) The different panels exhibit GFP fluorescence in the region infiltrated with ORF-IV at 3, 5 and 7 days post infiltration (dpi). The region infiltrated with empty vector (EV) served as control. (**B**) Northern blots of GFP transcript and GFP sRNA in regions infiltrated with ORF-IV (OIV) and EV (**C**) The graph represents the normalized Integrated Density Values (IDV) for the GFP and ORF-IV transcripts relative to 18S control cDNA at 3dpi (d3), 5dpi (d5) and 7dpi (d7) (**D**) The graph represents the normalized values of GFP transcripts and GFP siRNA as measured in the infiltrated regions at 3 dpi and 5dpi.

The suppressor activity was confirmed by validating the levels of GFP transcript and GFP small RNAs using northern blot (**Fig. 2B**). It was seen that GFP transcripts accumulated only in the regions where ORF-IV was transcribed indicating its interference with the silencing machinery. The time kinetics revealed that GFP transcript started forming by 3dpi even though significant florescence levels could be detected at 5dpi and 7dpi (**Fig. 2C**). Correspondingly, very low levels of GFP small RNAs were observed in leaf regions infiltrated with ORF-IV while prominent levels of the GFP small RNAs were detected in the empty vector infiltrated leaf sections (**Fig. 2B,D**). The negative correlation between the small RNA and ORF-IV transcript accumulation confirmed the suppressor action of the protein.

### 3. RTSV protein enhances the suppressor activity of ORF-IV

It is well known that Tungro disease is manifested by the co-infection of RTBV and RTSV. The presence of RTBV alone causes mild disease symptoms, but these are accentuated in the presence of RTSV (Hull, 1996; Borah et al., 2013). Thus it was important to further investigate the nature of RTBV ORF-IV RNA silencing suppression activity in presence of RTSV. For this the same assay system was employed and the leaves were infiltrated respectively with ORF-I, and ORF-II of RTBV and RTSV ORF encoding protease, coat protein 3 and polymerase as co-infiltrations with the ORF-IV (**Fig. 3)**. The infiltrations with ORF-IV alone served as reference point for assaying the suppressor activity.

**Figure 3.**
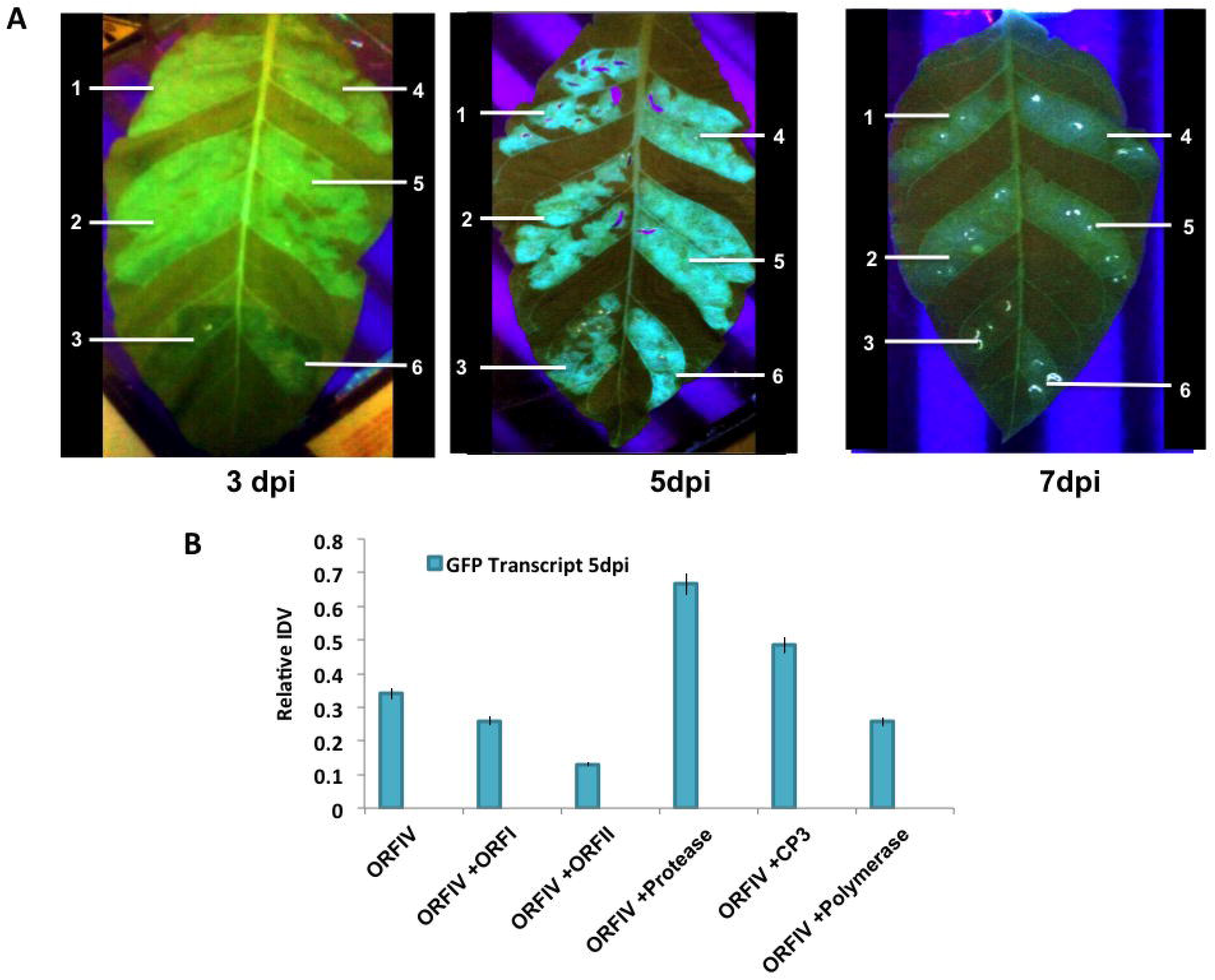
Reversal of GFP silencing by co-infiltration of RTBV and RTSV ORFs. (**A**) The different panels exhibit GFP fluorescence in the infiltrated regions at 3, 5 and 7 days post infiltration (dpi). The markings indicate region infiltrated with (1) ORF-IV construct, (2) ORF-IV co-infiltrated with RTBV-ORF-I, (3) ORF-IV co-infiltrated with RTBV ORF-II, (4) ORF-IV co-infiltrated with RTSV ORF coding for coat protein 3, (5) ORF-IV co-infiltrated with RTSV ORF coding for protease and (6) ORF-IV co-infiltrated with RTSV ORF coding for polymerase. (**B**) Normalized values for GFP transcripts in the different infiltrated regions at 5 dpi, confirmed using RT PCR.

It was observed that at 3dpi, weak suppressor activity is observed in regions co-infiltrated with ORF-IV and RTSV ORF encoding coat protein 3 (CP3) and polymerase. At 5dpi the suppression activity was significantly enhanced in individual co-infiltrations with RTSV ORFs but decreased in presence of the RTBV ORF-II. The enhanced suppression activities in regions co-infiltrated with RTBV ORF-IV and RTSV ORFs encoding protease and CP3 were sustained up to 7dpi (**Fig. 3A**). To validate the results, molecular analysis of the co-infiltrated zones was performed for GFP as well as the ORF transcripts using RT-PCR. The relative band intensity density values were calculated and normalized with respect to 18S control (**Fig. 3B**). Higher level of expression was seen for GFP in the regions where ORF-IV was co-infiltrated with RTSV ORFs coding for protease and CP3, indicating their role in enhancement of suppression activity encoded by RTBV. This clearly demonstrates that the early onset of suppression of ORF-IV is enhanced and sustained by the activity of RTSV ORFs, which may potentially lead to a potent infection in the rice plant. It was also observed that the respective presence of RTSV protease and CP3 caused a transient spread of silencing beyond the infiltrated zone (**Fig. 3A**). This indicates the suppression of both the local siRNAs as well as the mobile signals responsible for the systemic spread of silencing, however further investigation is required to understand the mechanism of action.

### 4. RTBV ORF-IV interacts with RTSV CP3 protein

To confirm if enhancement in ORF-IV suppressor activity by RTSV proteins is due to direct interaction, yeast two hybrid (Y2H) assay was performed. Yeast cells co-expressing binding domain (BD) fused with ORF-IV and activation domain (AD) fused with CP3 could grow on both three-drop-out media (without amino acids: leucine, tryptophan, and histidine) and four-drop-out media (without amino acids: leucine, tryptophan, histidine and adenine). The results show a direct and strong interaction of RTBV ORF-IV with RTSV CP3 (**Fig. 4** and **Fig S2**). The other proteins of RTSV coding for CP1, CP2, P1, polymerase and protease did not show any interaction with RTBV ORF-IV.

**Figure 4.**
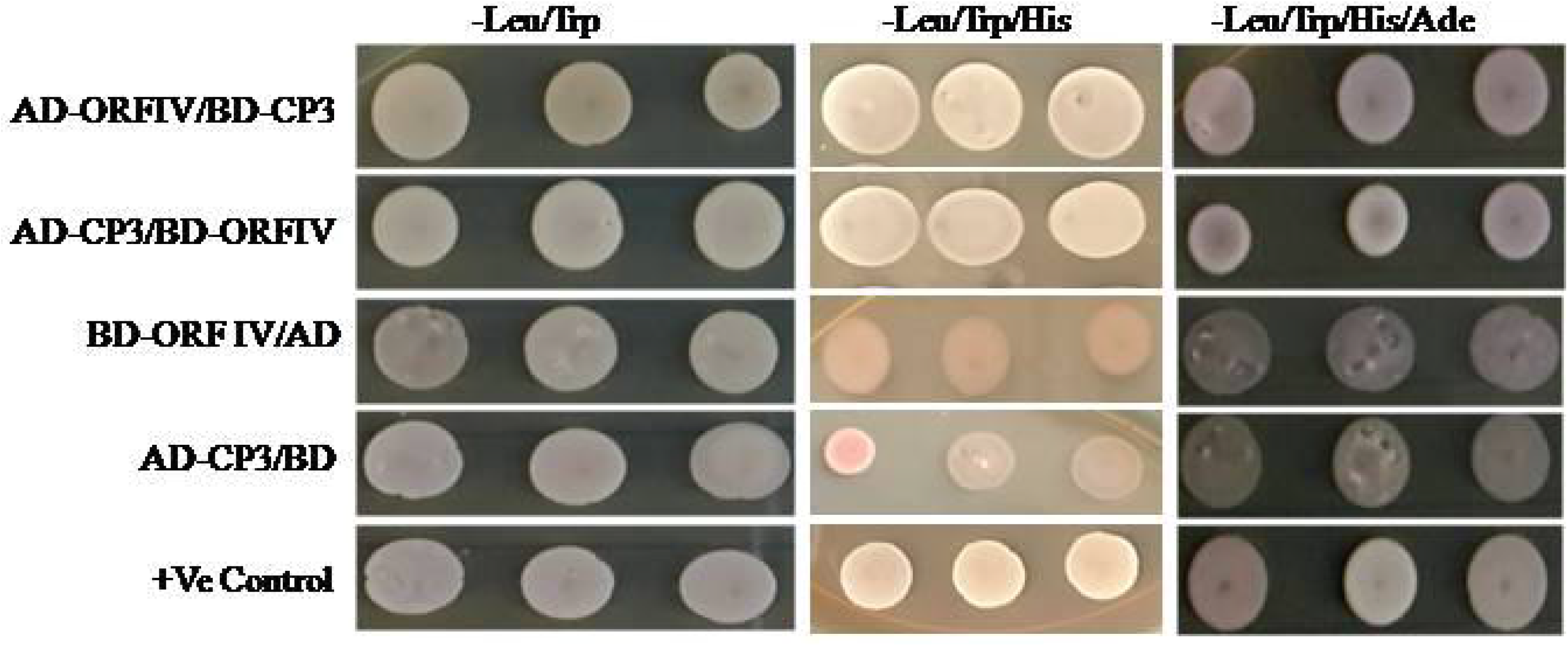
Yeast two-hybrid assay for direct interaction study. Yeast colonies of pGAD-CP3 and pGBD-ORF-IV were co-transformed and selected on (i) two drop out plate (Leu^−^ Trp^−^), (ii) triple drop out (Leu^−^ Trp^−^ His^−^) (ii) triple drop out (Leu^−^ Trp^−^ His^−^) supplemented with 5 μ M 3-AT. Co-infiltration of pGAD-CP3 with pGBD and pGAD with pGBD-ORF-IV was performed as control.

## Discussion

The host RNA silencing mechanisms are triggered in response to viral infection to produce siRNAs as a first line of defense (Llave, 2010; Pantaleo, 2011; Rajeswaran and Pooggin, 2012; Szittya and Burgyán, 2013). The siRNA molecules can potentially act by directing the cleavage of the complementary viral transcripts or by inducing transcriptional silencing. During Tungro virus disease, the full-length RTBV transcript can act as the template for reverse transcription-mediated replication of the viral DNA (Hull, 2013), so it appears to be the most suited inducer of RNA silencing. The precursors of viral siRNAs are possibly produced by Pol II-mediated bi-directional transcription of the viral DNA (Aregger et al., 2012; Rajeswaran et al., 2012). The viral double stranded RNAs can be processed by DCL2 and DCL3 to generate the 22-nt and 24-nt siRNAs, respectively whereas DCL1 and DCL4 process 21-nt siRNAs (Blevins et al., 2006). It has been shown that 21-nt siRNAs associate with AGO1 proteins to direct cleavage of target mRNAs and control the local spread of silencing (Hamilton et al., 2002; Himber et al., 2003; Dunoyer et al., 2010; Molnar et al., 2010). While 24-nt siRNAs associate with AGO4 to direct cytosine methylation of complementary target DNA (Song et al., 2012; Wu et al., 2010; Wei et al., 2014) and are responsible for the systemic spread of silencing.

Analysis of the small RNA libraries prepared from tungro-infected leaves identified the 21-nt small RNAs as the largest and most diverse group. The 21-nt species are normally associated with post-transcriptional gene silencing for allowing an immediate recognition and elimination of incoming viruses (Hamilton et al., 2002). Almost 50% of the small RNA reads aligned to the ORF-IV region, although major clustering was towards 3’ end. The ORF-IV transcript arises by splicing and joining the coding sequences to the 5’ untranslated region (Fütterer et al., 1994). It has been demonstrated that, in DNA viruses the intergenic regions harboring bidirectional promoter elements between the transcription start sites are a poor source of viral siRNAs (Wang et al., 2011; Aregger et al., 2012) while the transcripts of either polarity are preferentially sources of double strand RNA, that are subsequently processed by the DCLs (Blevins et al., 2011).

The *in planta* silencing suppression assay was based on screening for reversal of GFP silencing in stably silenced tobacco plants. High levels of transgenic RNA are recognized as aberrant by the rate limiting cellular cofactors and this triggers its conversion into a duplex by the host RNA dependent RNA polymerase (Takeda et al., 2002). The dsRNA initiates siRNA production, which in turn leads to cleavage of transcripts called sense PTGS (S-PTGS). Both DCL4- and DCL2- processed secondary siRNAs were reported to be effective in silencing activity when an inducing transgene is expressed constitutively (Fusaro et al., 2006; Mallory and Vaucheret, 2006; Moissiard et al., 2007). At later stages the siRNAs can guide the protein complexes to switch off gene expression, thereby establishing the stable silenced state for the gene (Agrawal et al., 2003). The theory is substantiated by the frequent induction of silencing by over-expression of the sense gene (Napoli et al., 1990; Di Serio et al., 2001). The transgene-induced and virus-induced gene silencing overlap mechanistically, so the process has been exploited to identify the suppressors of RNA silencing (Karjee et al., 2008) by its action of reversing the inhibition of GFP expression.

The ORF-IV demonstrated RNA silencing suppression activity *in planta*. It has been reported that RTBV protein encoded by ORF-IV may not be essential for viral replication, assembly or movement as other pararetroviruses belonging to the family Caulimoviridae do not possess any related gene. This locus is a hot spot for siRNA generation so it is likely to have evolved the ability to counteract the host RNA silencing defenses (Zvereva and Pooggin, 2012). An indication for this was provided by GFP co-infiltration experiments wherein ORF-IV was shown to suppresses the silencing signal in boundaries of infiltrated zones, even though it was reported to enhance cell autonomous silencing (Rajeswaran et al., 2014). However, this study identified ORF-IV as a weak suppressor of the S-PTGS or pre-established stable silencing and the results were supported by molecular validations.

This study demonstrates that the RTSV proteins may help in early onset of suppression by ORF-IV and sustenance of the activity, which potentially leads to a potent infection in the rice plant. It should be noted that both RTBV and RTSV are required to produce typical viral symptoms of tungro disease (Hull, 1996). The results of co-infiltration experiments showed that suppression activity of ORF-IV was enhanced in presence of RTSV ORF-protease and CP3. It was also observed that the presence of RTSV protease and CP3 caused a transient spread of silencing beyond the infiltrated zone. Notably, RNAi based silencing of CP3 gene caused restrain of tungor disease in rice (Malathi et al 2019). Important role of RTBV ORF-IV and RTSV CP3 in virus diversification and evolution has been suggested (Mangrauthia et al., 2010; Mangrauthia et al., 2012). Yeast based interaction studies indicated direct interaction between RTSV and RTBV proteins. It is reported that RTSV individual infection does not cause any major disease symptoms but in presence of RTBV the severity of infection increases. RTSV is mainly known to promote the transmission of the disease (Hibino et al., 1978).

The results obtained in this study give a clear indication on the complexity of Tungro disease driven by the synergy of two disparate viruses. The study brings to focus the important role of RTBV ORF-IV in disease manifestation. This locus acts as both, victim and silencer of the RNA silencing pathway in host plants representing a key target that can be exploited for boosting host plant resistance against tungro virus infection. The study also provides evidence on the complementation of RNA suppression activity of RTBV (ORF-IV) by RTSV (CP3) and the direct interaction between the two proteins. This implies at the necessity of presence of both the viruses to cause the disease. It can thus be hypothesized that RTBV encodes suppression activity for handling the localized silencing activity of the host plant whereas the RTSV components help in suppressing the cell to cell spread of silencing, thus sustaining the spread of infection. The silent role of RTSV CP3 to enhance and extend the suppression activity of ORF-IV indicates that nature and degree of interaction of the viral proteins with each other in the host need to be further elucidated to understand the complexity of Tungro disease manifestation and its protraction.

## Acknowledgements

The research was supported by financial grants received from the Department of Biotechnology, Government of India. AA acknowledges the fellowship received from CSIR, India.

## Compliance with Ethical Standards

N.A.

## Ethical approval

This article does not contain any studies with human participants or animals performed by any of the authors.

## Authors Contributions

AA and NSM established the overall concept and performed the analysis. AT wrote the PERL scripts for the analysis. MP and SKM cloned the viral ORFs and performed the interaction studies. NSM and AA prepared the manuscript. All the authors read and approved the final manuscript.

## Supplementary data

**Figure S1. Genomic organization and structure of RTBV and RTSV.**

**Figure S2. Plates showing the yeast two-hybrid interaction of pGAD-CP3 and pGBD-ORF-IV.** Yeast colonies were co-transformed and selected on (A) two drop out medium (Leu^−^ Trp^−^) and (ii) four drop out medium (Leu^−^ Trp^−^ His^−^, Ade^−^). Co-infiltration of pGAD-CP3 with pGBD and pGAD with pGBD-ORF-IV was performed as control. The positive control was same as that provided in the kit. For negative control empty AD and BD vectors were co-transformed.

