## Supplementary figures and images for "Coordinated action of RTBV and RTSV proteins suppress host RNA silencing machinery"

### Supplementary File 1

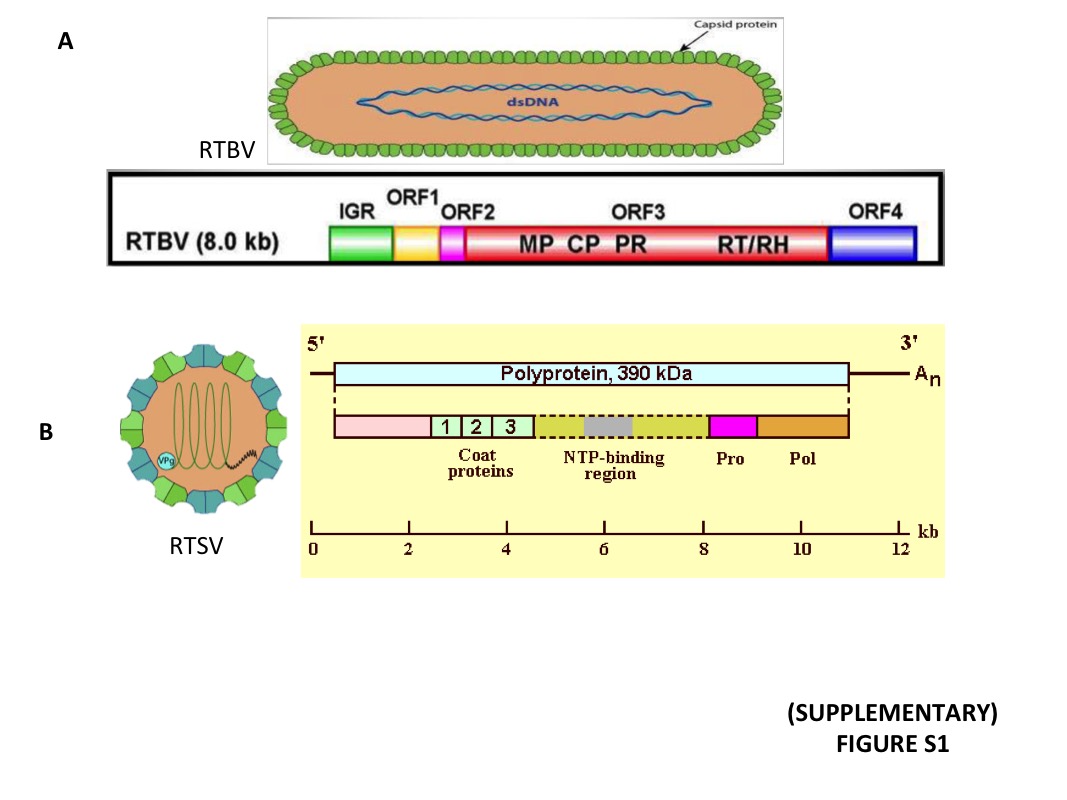

### Supplementary File 1

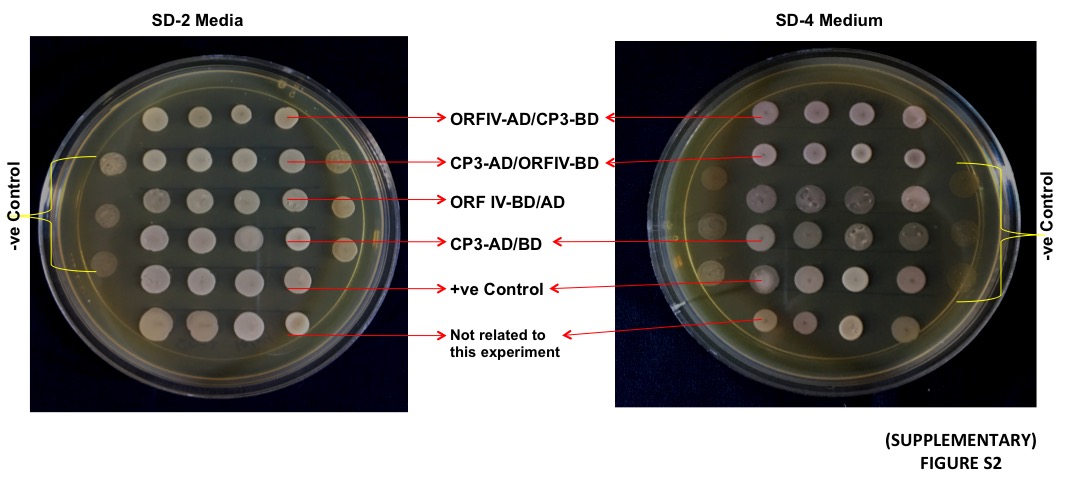
